# Age-related Structural and Functional Variations in 5967 Individuals across the Adult Lifespan

**DOI:** 10.1101/810101

**Authors:** Na Luo, Jing Sui, Anees Abrol, Dongdong Lin, Jiayu Chen, Victor M. Vergara, Zening Fu, Yuhui Du, Eswar Damaraju, Yong Xu, Jessica A. Turner, Vince D. Calhoun

## Abstract

Exploring brain changes across the human lifespan is becoming an important topic in neuroscience. Though there are multiple studies which investigated the relationship between age and brain imaging data, the results are heterogeneous due to small sample sizes and relatively narrow age ranges. Here, based on year-wise estimation of 5967 subjects from 13 to 72 years old, we aimed to provide a more precise description of adult lifespan variation trajectories of gray matter volume (GMV), structural network correlation (SNC) and functional network connectivity (FNC) using independent component analysis and multivariate linear regression model. Our results revealed the following relationships: 1)GMV linearly declined with age in most regions, while parahippocampus showed an inverted U-shape quadratic relationship with age; SNC presented a U-shape quadratic relationship with age within cerebellum, and inverted U-shape relationship primarily in the default mode network (DMN) and frontoparietal (FP) related correlation. 2) FNC tended to linearly decrease within resting-state networks (RSNs), especially in visual network and DMN. Early increase was revealed between RSNs, primarily in FP and DMN, which experienced decrease at older ages. U-shape relationship was also revealed to compensate for the cognition deficit in attention and subcortical related connectivity at late years. 3) The link between middle occipital gyrus and insula, as well as precuneus and cerebellum, exhibited similar changing trends between SNC and FNC across the adult lifespan. Collectively, these results highlight the benefit of lifespan study and provide a precise description of age-related regional variation and SNC/FNC changes based on a large dataset.

## INTRODUCTION

Human lifespan development is a major topic of interest in neuroscience. The brain maturation process is likely to experience a predominantly genetically determined growth first, followed by a more plastic gene-environment interaction period (Cao, Huang, & He, 2017; Collin & van den Heuvel, 2013; van den Heuvel et al., 2013). Neurodevelopmental trajectories in gray matter (GM) volumetric variations have been extensively studied (Anders M. Fjell, McEvoy, Holland, Dale, & Walhovd, 2014; Kennedy et al., 2009; Narvacan, Treit, Camicioli, Martin, & Beaulieu, 2017; Terribilli et al., 2011). Gray matter (GM) reduction of normal aging in frontal, temporal, parietal and insular area were largely reported (Abe et al., 2008; Farokhian, Yang, Beheshti, Matsuda, & Wu, 2017; Raz & Rodrigue, 2006). However, conflicting results are also observed when using different sample sizes and age ranges; one such area is the hippocampus. Linear relationship was revealed when evaluating only in older ages, while nonlinear variations of this regions were observed when the samples covered a long lifespan from teenager to older ages (J. S. Allen, Bruss, Brown, & Damasio, 2005; A. M. Fjell et al., 2013).

Cerebral alterations were characterized by the reorganization of cortical connectivity patterns (Meunier, Achard, Morcom, & Bullmore, 2009; Wu et al., 2012). Several previous studies have demonstrated that patterns of structural connectivity or functional connectivity undergo characteristic variations across the lifespan (Cao et al., 2014; Wang, Su, Shen, & Hu, 2012; Yang et al., 2014; Zuo et al., 2010). For example, a numbers of studies focused on changes in functional network connectivity (FNC) within and between networks, have stated that FNC tended to decrease within resting-state networks (RSNs) with aging, including the visual network and default mode network (DMN) (E. A. Allen et al., 2011), and increase between RSNs, especially between components of the somatomotor network, the ventral attention network and dorsal attention network, which were best fit by convex quadratic models (Betzel et al., 2014). Functional network connectivity (FNC) can be extracted from different methodologies, including independent component analysis (ICA) and ROI-based network construction. In this study, FNC matrix were estimated by computing the correlation among pairs of time courses identified from ICA as (Calhoun, Adali, Pearlson, & Pekar, 2001; Jafri, Pearlson, Stevens, & Calhoun, 2008; Segall et al., 2012; Xu, Groth, Pearlson, Schretlen, & Calhoun, 2009), which capture networks that covary across time courses. In parallel, we followed our previous paper (Erhardt, Allen, Damaraju, & Calhoun, 2011; Segall et al., 2012) to define the relationships between different structural ICA derived components as structural network correlations (SNC).

Overall, the present study aimed to replicate previous studies and to provide a more robust and precise elaboration of age-related variations using a much larger sample size (5967 subjects), and due to the plenty of sample size at every age year from 13 to 72, we are able to investigate the changing trend of SNC and FNC year-wisely, using ICA and a multivariate linear regression model (MLR). Moreover, based on a well-matched structural-functional template, we compared how age-related FNC changes similar to age-related SNC variations. We performed 3 analyses in response to the following hypotheses regarding the age-varying imaging discoveries: (1) As for the GM volume, we expected hippocampus or para-hippocampus would show a close association with age increase, since this region has been widely reported to be sensitive to aging (Bartsch & Wulff, 2015; Burke et al., 2018). (2) For structural or functional network investigation, we hypothesize that quadratic relationships would be revealed between FNC or SNC of brain regions in charge of higher-order cognitive processing, since the late-maturing brain regions are revealed to be more sensitive to the deleterious effects of aging (Kalpouzos et al., 2009; Toga, Thompson, Mori, Amunts, & Zilles, 2006; Zuo et al., 2017). (3) An exploratory analysis: after examining the age-varying FNC and SNC patterns, we compared between each other and expected to find certain similarity between functional and structural links.

## MATERIALS AND METHODS

### Data acquisition and preprocessing

The data used in this study consisted of 6101 structural magnetic resonance imaging (MRI) scans and 7500 resting-state functional MRI (fMRI) scans, which were collected at the University of New Mexico (UNM) and the University of Colorado Boulder (UC, Boulder). Data in the UC, Boulder site was collected using a 3T Siemens TIM Trio MRI scanner with 12 channel radio frequency coils, while data in UNM site was acquired using the same type of 3T Siemens TIM Trio MRI scanner, and a 1.5T Avanto MRI scanner. The site effect will be controlled in the subsequent analysis. All the data were previously collected, anonymized, and had informed consent received from subjects. As the data is a de-identified convenience dataset, we do not have access to the health and identifier information and this dataset likely includes some individuals with brain disorders. We have confirmed that the brain images do not have any obvious pathology or atrophy.

T1-weighted structural images were acquired with a five-echo MPRAGE sequence with TE = 1.64, 3.5, 5.36, 7.22, 9.08 ms, TI = 1.2 s, TR = 2.53 s, flip angle = 7°, number of excitations = 1, field of view = 256 mm, slice thickness = 1 mm, resolution = 256 × 256. The structural data were preprocessed based on voxel-based morphometry (VBM) in SPM12(Ashburner & Friston, 2005). The preprocessing pipeline includes: (1) spatial registration to a reference brain; (2) joint bias correction and tissue classification into gray matter, white matter and cerebrospinal fluid using SPM12 old segmentation; (3) spatial normalization to the standard Montreal Neurological Institute (MNI) space using nonlinear transformation; (4) modulation by scaling with the amount of volume changes, and (5) smoothing to 10×10×10 mm FWHM (Matt Silver, Montana, Nichols, & Alzheimer’s Disease Neuroimaging, 2011; Sui et al., 2013). The smoothed gray matter volume (GMV) images from each dataset were spatially correlated to the mean image to assess outliers. Scans with a correlation less than 0.7 were removed.

The fMRI images were used in a previous study that evaluated replicability in time-varying functional connectivity patterns (Abrol et al., 2017), which has clearly reported the acquisition parameters and preprocessing pipelines. T2*-weighted functional images were acquired using a gradient-echo EPI sequence with TE = 29 ms, TR = 2 s, slice thickness = 3.5 mm, flip angle = 75°, slice gap = 1.05 mm, matrix size = 64 × 64, field of view = 240 mm, voxel size = 3.75 mm × 3.75 mm × 4.55 mm. The data preprocessing pipeline included discard of the first three images for the magnetization equilibrium, realignment using INRIalign (Freire & Mangin, 2001), timing correction with the middle slice as reference, spatial normalization into the MNI space. Images collected at 3.75 mm × 3.75 mm × 4.55 mm were then slightly upsampled to 3 mm × 3 mm × 3 mm, resulting in a data cube of 53 × 63 × 46 voxels. The upsampled images were further smoothing with a 10 mm Gaussian model (M. Silver, Montana, Nichols, & Neuroimaging, 2011). The fMRI data covered the entire cerebellum. Anomaly detection in the form of a correlation analysis on the five upper and lower slices of the functional images was performed on all 7500 scans in order to detect scans that failed the reorientation process, or had any missing slices. This outlier detection removed 396 subjects, thus leaving behind a total number of 7104 subjects corresponding to approximately 95% of the available data. The time courses for all subjects were post-processed in the FNC construction step to remove any residual noise sources.

After preprocessing, 5967 scans were retained with both structural and functional MRI images. The complete demographic information were shown in Table 1.

**Table 1.**
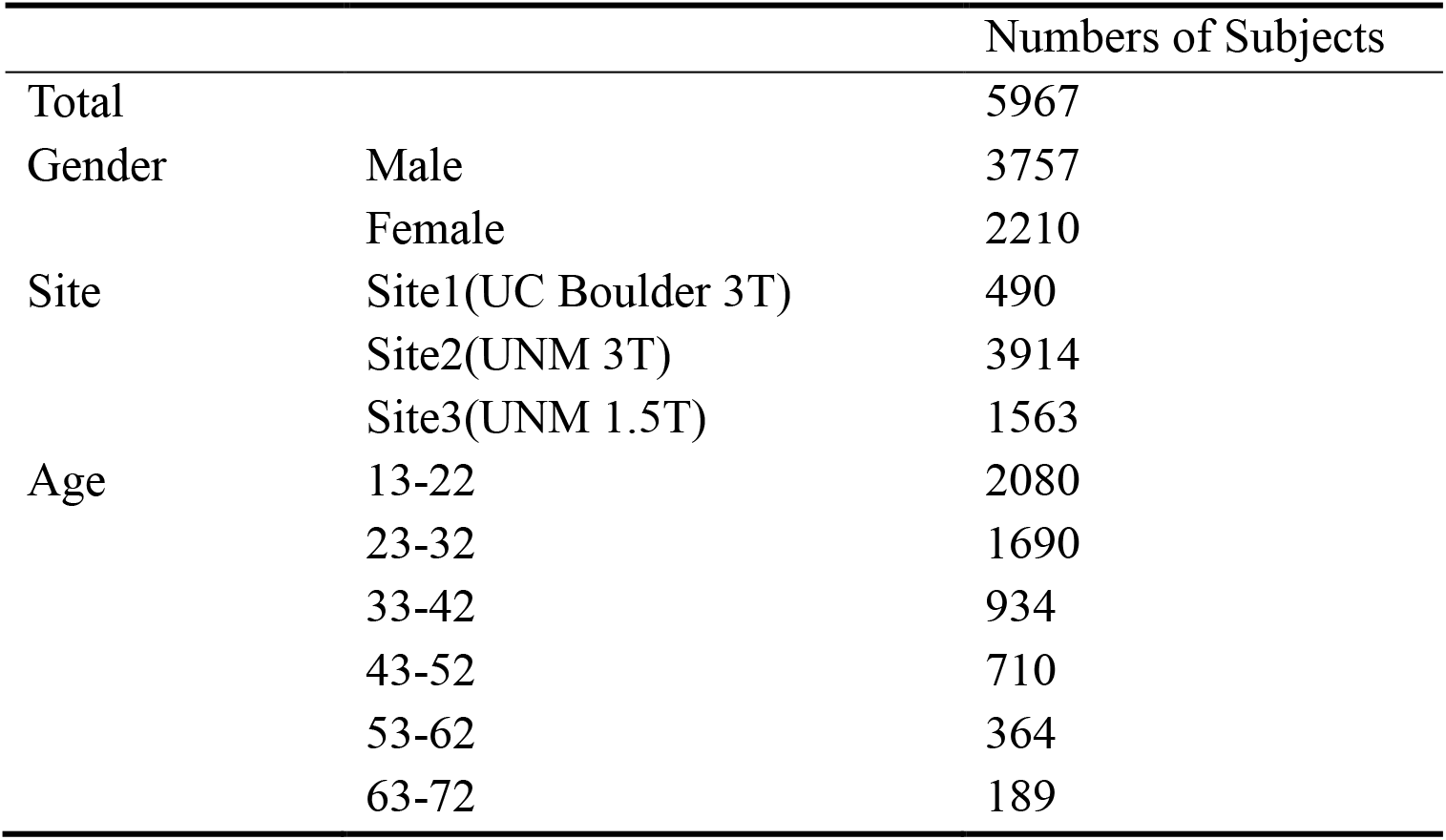
Demographic information

### Data statement

The structural and functional data used in the present study can be accessed upon request to the corresponding authors.

### Independent components derived from ICA for functional and structural data

ICA analysis on the functional data was conducted in our previous study (Abrol et al., 2017) using group ICA (GICA) implemented in the GIFT toolbox (http://mialab.mrn.org/software/gift/) (Calhoun, Adali, Pearlson, & Pekar, 2002). The number of components was set to be 100. After visual inspection of all the 100 components, 61 components were selected with peak activations in gray matter, time courses dominated by low-frequency fluctuations, and high spatial overlap with resting networks. We then defined nine domains/networks based on Yeo *et al.*’s seven-network template (Yeo et al., 2011), which added cerebellar and subcortical network as another two networks: visual network (VIS), somatomotor network (SM), dorsal attention network (DA), ventral attention network (VA), subcortical network (SUB), limbic network (LIMBIC), frontoparietal network (FP), DMN and cerebellar network (CB). We further sorted the 61 components into the nine domains for further analysis as shown in **Figure S1**. The criteria for sorting the components was based on the peak location.

ICA decomposition on the structural data were investigated with source based morphometry (SBM), which decomposed the GMV images into a loading parameter matrix (the A matrix in Figure 1A) and a source matrix (the S matrix in Figure 1A) (Calhoun et al., 2001; Xu et al., 2009). The loading parameters (LP) matrix represented the weight of components for each subject and the source matrix indicated the corresponding spatial maps. For the purpose of comparing the similarities between age-related structural and functional changes, we used the same numbers of components (100) as functional data for ICA analysis. Components with significant spatial overlap with ventricles, large vasculature, white matter and the brainstem, or located at the boundaries between these regions and GM were excluded as (Du et al., 2015). Of all the 100 structural components identified from ICA, 71 GM components were retained for analysis after removal of artifact components via visual inspections and further divided into eight domains as defined by (Yeo et al., 2011) (**Figure S2**).

**Figure 1.**
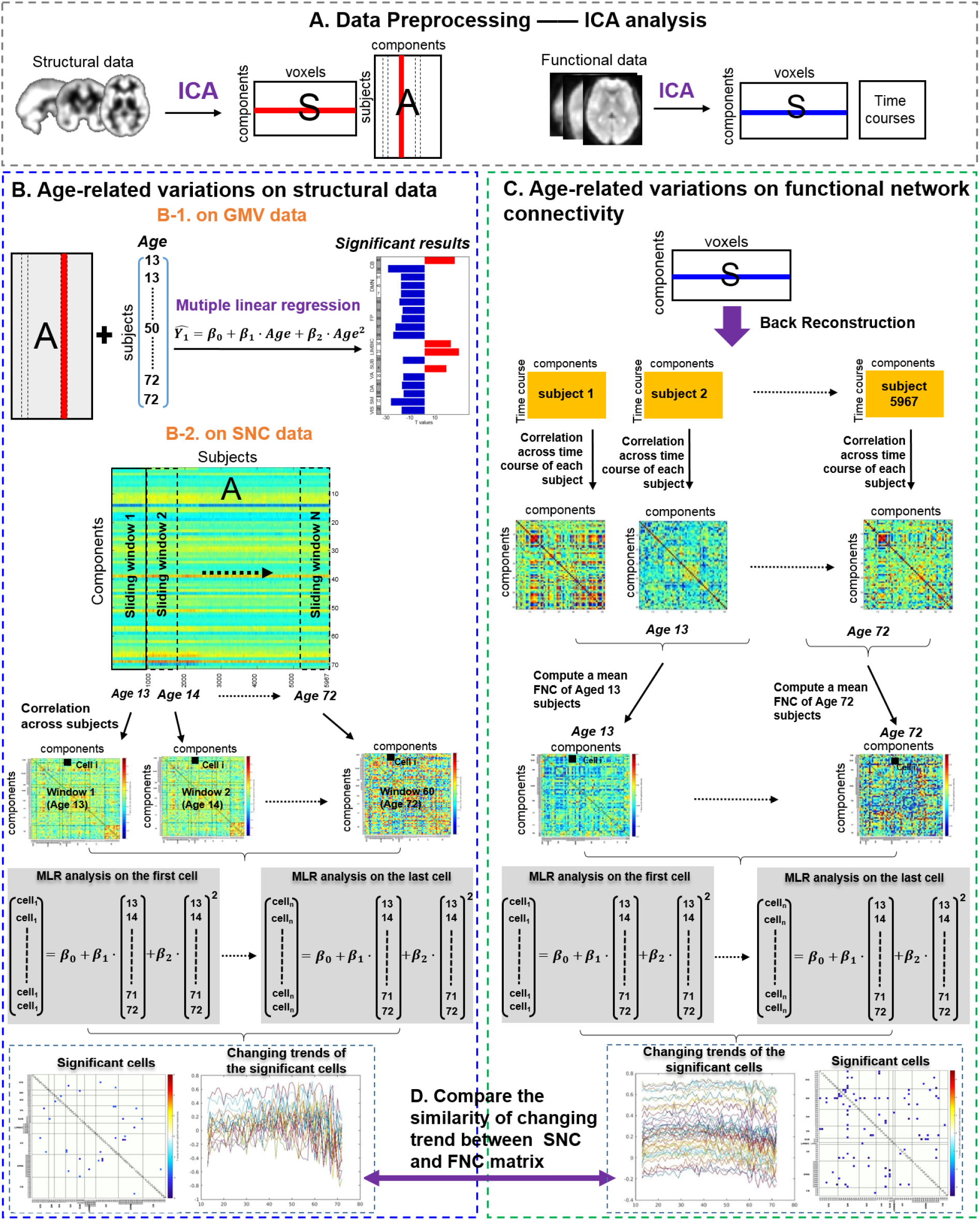
Illustration of the analysis pipeline. **A)** Structural and functional data were first decomposed by ICA. **B)** The analysis pipeline of computing the relationship between age and structural data. B-1: We first computed the relationship between each component in the A matrix and age using multiple regression model (MLR) model, which measures how GMV changes across the adult lifespan. B-2: We then applied a sliding-age window to the structural loading parameters [A matrix] to construct structural network correlation (SNC) matrix for each age stage and further examined the relationship between age and SNC using MLR. **C)** For the functional data, group-level spatial maps [S] were used to back reconstruct the matrix [time courses by components] for each subject and then functional network connectivity (FNC) was constructed cross time courses for each subject. We further sorted the subjects in an increasing order to compute the mean FNC matrix for each age stage and examined the relationships between age and FNC using the same MLR model. **D)** We finally measured the similarity of the changing trends for the significant cells between SNC and FNC based on a structural-functional matched template.

### Construction of age-resolved structural and functional network connectivity

#### Sliding window network construction from structural data

We constructed the SNC matrix by computing the correlations among the structural loading parameters, which measures the links between pairs of covarying gray matter patterns identified from SBM analysis (Segall et al., 2012). Age-related SNC matrices were constructed from 13 to 72 years old using a sliding window method (Figure 1B-2). Loading parameters were cross-correlated within windows that contained participants of the same age and incrementally moved across the age-range in regular increments (Vasa et al., 2018). The step size was set by 1 year old. The window width depended on the number of subjects in each age stage. A partial correlation, using gender, site, and age × gender as covariates, was used to compute the SNC, then 60 SNC matrices would be constructed corresponding to the 60 age stages.

#### Network construction from functional data

Back reconstruction using group information guided ICA (GIG-ICA) was performed after GICA, with the selected 61 components as reference, to estimate subject-level time courses and maps for each subject (Du & Fan, 2013). The back-reconstructed component time-courses went through additional processing steps to remove any residual noise sources mostly including low frequency trends originating from the scanner drift, motion related variance emerging from spatial non-stationarity caused by movement, and other non-specific noise artifacts unsatisfactorily decomposed by the implemented linear mixed model. More specifically, the post-processing steps featured de-trending existing linear, quadratic and cubic trends, multiple linear regression of all realignment parameters together with their temporal derivatives, outlier detection using 3D spike removal, and low pass filtering with high-frequency cut-off being set to 0.15 Hz. Lastly, the time courses were variance normalized. An FNC matrix was finally constructed across time courses for each subject with gender, site and age × gender as covariates (Jafri et al., 2008). Subjects were further sorted in an increasing order, and we computed a mean FNC matrix for all FNC matrices in the same age stage (from 13 to 72 years old) (Figure 1D). Accordingly, 60 FNC matrices were constructed corresponding to 60 age stages.

### Correlations between age and GMV, SNC and FNC

After ICA decomposition, we first computed the relationship between structural loading parameters and age using MLR model as shown in Figure 1B-1, which measures how GMV changes across the adult lifespan. Then the age-related variation of connectivity/correlation in the constructed SNC matrix and FNC matric were also measured using MLR model as shown in Figure 1B-2 and **1C**. Here, we model *Ŷ* as the variable of interest, which represents the structural loading parameters, or the value across all windows for each cell in the SNC/FNC matrix. Then we applied the following MLR models:

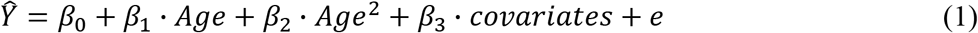

where, *Ŷ* is a [5967×1] vector for the GMV case and a [60×1] vector for the SNC/FNC case. *Age* represents a matrix that includes ages with associated parameters *β*_1_. *Age*^2^ is a matrix consisting of squared ages with the associated parameter *β*_2_, and *e* is an error term. When computing the relationship between structural loading parameters and age, covariates indicated gender, site and age × gender. As these covariates were already regressed when constructing the SNC and FNC matrices, the *covariates* term was crossed out when measuring the relationship between SNC, FNC matrices and age. Cells with significant p-values (FDR<0.05) of parameter *β*_1_ indicated an age-related linear relationship, and cells with significant p-values (FDR<0.05) of parameter *β*_2_ indicated an age-related quadratic relationship. As 71 structural components were used to measure the relationship between structural loading parameters and age using the MLR model, FDR correction for the significance of *β*_1_ and *β*_2_ were based on the 71 p-values separately. When used the MLR model to select the significant cell in the SNC matrices across 60 age stages, 2485 cells ((71 × 70)/2) were computed to select the significant cells. For FNC, 1830 cells ((61×60)/2) from each FNC matrix were measured to select the significant cells. At last, T maps corresponding to these significant cells were further plotted.

### Comparisons between age-related SNC and FNC variations

To further compare similarities between age-related SNC changes and FNC variations, we adopted a matched structure-function template revealed in our current under-review study **(Figure S3)**. Based on the template, we could find the matching cells between SNC and FNC matrix. Afterwards, we plotted curves with age of these matched cells and further computed the correlation between curves of each matched pair (Figure 1E).

## RESULTS

### Gray matter volume changes across the adult lifespan

Post hoc MLR analysis identified 19 components that exhibited a significant linear correlation with age (i.e., significance was measured using effect magnitude [partial R-square > 0.05] and FDR-corrected P values [p < 0.05]), and 15 components exhibited a linear declining relationship (Figure 2A). Components from cerebellar and somatomotor domains, especially the vermis and precentral area, showed the highest declining correlations with age (Figure 2B). Four other components (s-IC4, s-IC23, s-IC38 and s-IC49) exhibited a positive relationship with age, which were comprised of the thalamus, parahippocampal, hippocampus and parts of the cerebellum, respectively. The component that primarily consisted of the parahippocampus and temporal pole (s-IC38) further exhibited a quadratic relationship with age. By fitting the component into a quadratic plot, we estimated the turning point to be 43.68 years old as shown in Figure 2C.

**Figure 2.**
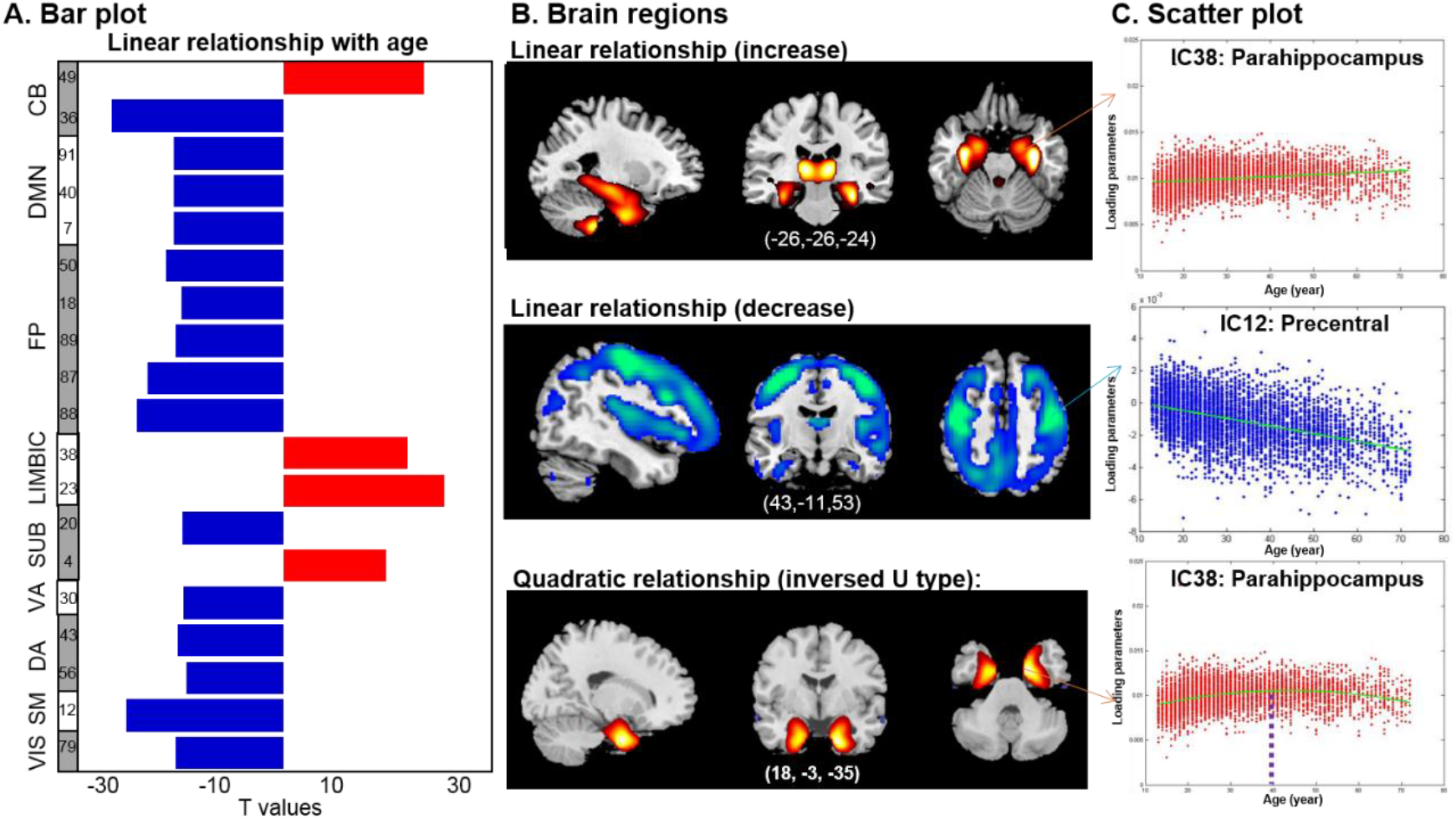
The relationships between age and ICA-decomposed components. A) The T values of the significant correlation between age and the structural components. B) The corresponding brain regions of the significant components; three types of relationship were revealed. C) The scatter plot of the representative components in each type.

### Structural network connectivity changes across the adult lifespan

Figure 3A and **3B** indicate the significant cells showing linear and quadratic SNC changes with age. No cells presented significant linear relationship with age, while quadratic relationship (FDR<0.05) were revealed, including both U-shape (13 cells) and inverted U-shape types (16 cells). To rule out the randomness, we further computed the pair-wise correlation between the 13 U-shape cells and 16 inverted U-shape cells. As shown in **Figure S4**, among all 208 pairs of cells, 96 pairs presented significant correlation after FDR correction (FDR<0.05). For each paired cells, we examined the relationship between age and one cell (U or inverted U-shape) while controlling for the other cell. As shown in **Table S1**, the majority of the significant age-cell quadratic relationship are retained (light blue background), suggesting that it is unlikely to identify both U and inverted U-shape relationship by randomness. Most significant cells with a U-shape relationship were observed within the cerebellar connectivity. The connectomes of the significant cells are plotted in Figure 3B-1. Figure 3B-2 describes the changing trends across the adult lifespan for all significant cells. By fitting these scatters into a quadratic plot, the turning point was estimated to be 45.17 years old. Meanwhile, inverted U-shape relationships were primarily revealed in DMN and FP related correlations. Figure 3B-3 and **3B-4** separately depict the connectome of the significant cells and the changing trends across the adult lifespan. By fitting these scatters into a quadratic plot, the turning point was estimated to be 40.83 years old.

**Figure 3.**
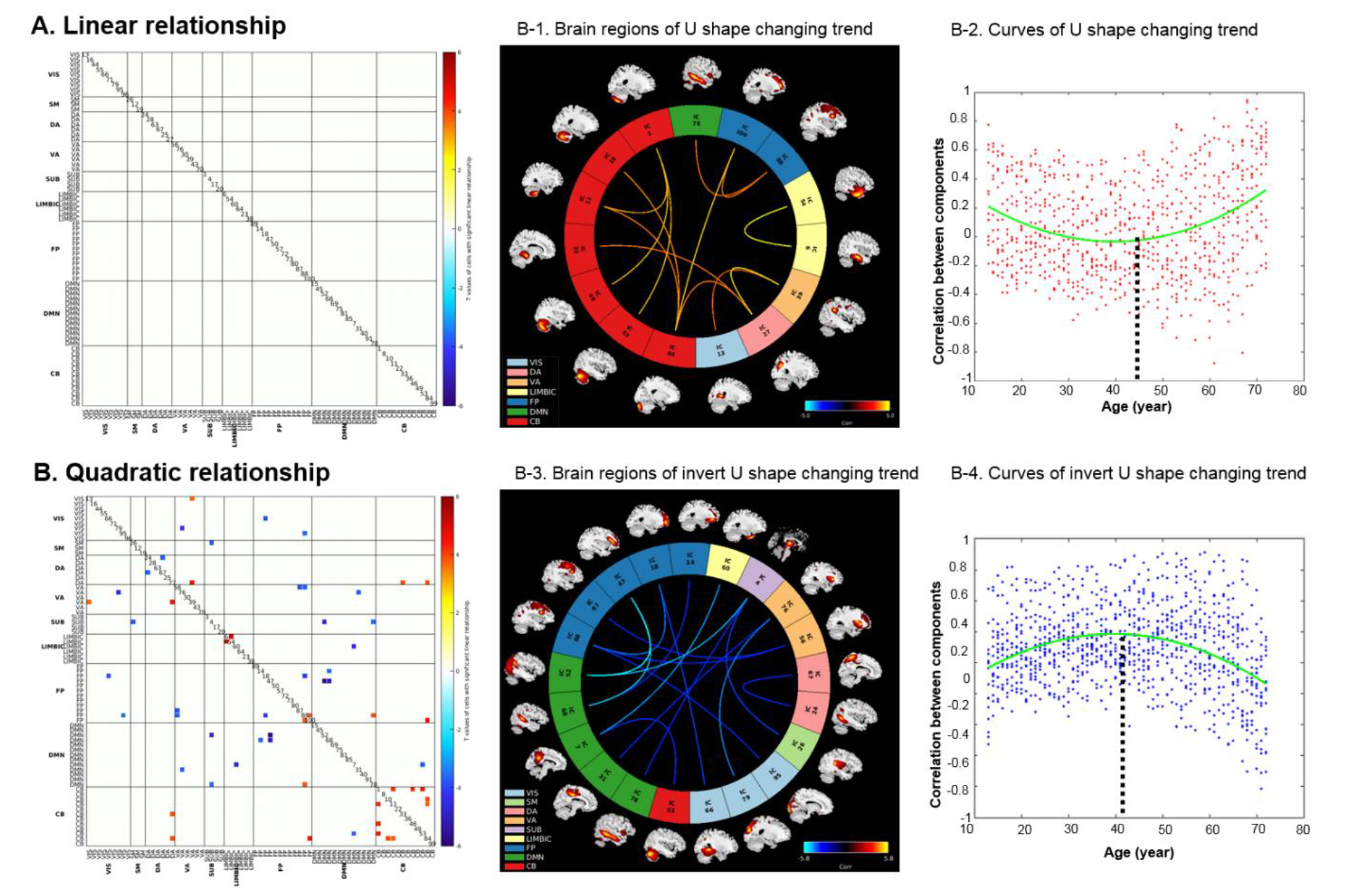
Correlations between age and SNC. A) No significant linear relationship was revealed. B) Both U-shape and inverted U-shape quadratic relationship were revealed. Figure 3B-1 and 3B-2 depict the connectome of the significant U-shape cells and the changing trends across the adult lifespan. Figure 3B-3 and 3B-4 separately depict the connectome of the significant inverted U-shape cells and the changing trends across the adult lifespan.

### Functional network connectivity changes across the adult lifespan

Figure 4A and **4B** present the significant cells showing positive and negative linear FNC changes with age separately (FDR<0.0001). The FNC within RSNs, especially the VIS and DMN domains, linearly decreased with age (Figure 4A). Linear increase was revealed between RSNs, primarily in FP and DMN (Figure 4B). The connectomes of the significant cells are plotted in Figure 4A-1 and Figure 4B-1. Figure 4A-2 and Figure 4B-2 describe the changing trends across the adult lifespan of all significant cells. U-shape relationship was primarily revealed in some VA and SUB related connectivity as shown in Figure 4C. Inverted U-shape relationships with age are revealed in Figure 4D (FDR<0.0001), centered on the regions like superior temporal gyrus (STG: rs-IC62), anterior cingulate gyrus (ACG: rs-IC94), middle cingulate gyrus (MCG: rs-IC67) and superior frontal gyrus (SFG: rs-IC100). The connectomes of the significant cells are plotted in Figure 4C-1 and **4D-1**. Figure 4C-2 and **4D-2** separately describe the changing trends of the significant cells from 13 years old to 72 years old. By fitting these scatters into a quadratic plot, the turning points were estimated to be 40 years old and 36.5 years old for U-shape and inverted U-shape relationships, respectively. To further examine the effect of head motion on our results, we first computed the mean framewise displacements (FD) for each subject (Power, Barnes, Snyder, Schlaggar, & Petersen, 2012; C. G. Yan et al., 2013). The relationship between mean FD and age was then examined, which indicated an un-significant result (*r*=0.0078, *p*=0.5485). Moreover, we added the mean FD as another covariate when computing the age-related significant FNC cells. As shown in **Figure S6**, the results are similar with the raw results, which suggests head motion doesn’t influence our results in this study.

**Figure 4.**
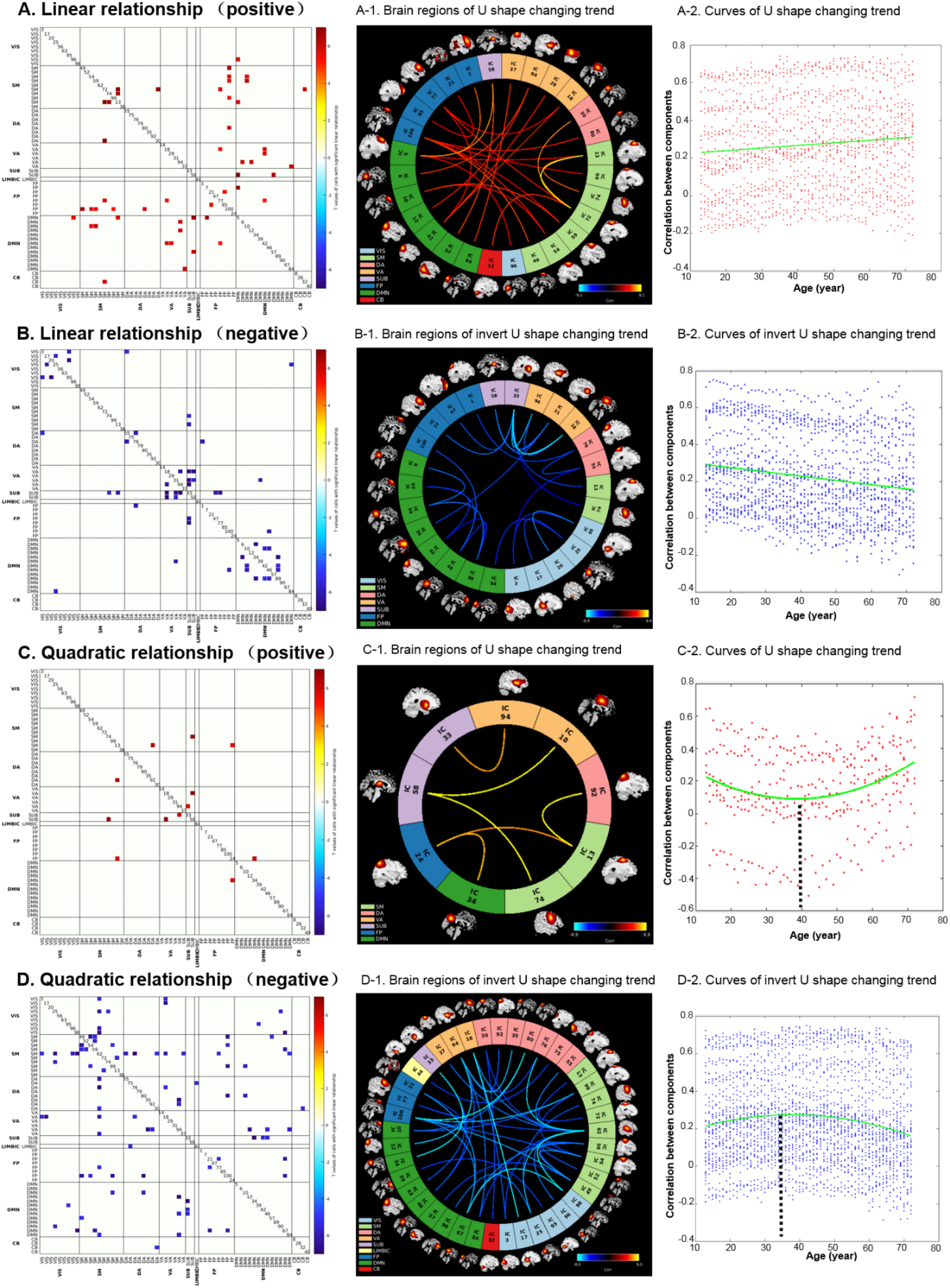
The relationship between age and FNC. A) Cells exhibiting a linear positive relationship with age; A-1: the connectome of the significant cells; A-2: the changing trends of the significant cells. B) Cells exhibiting a linear negative relationship with age; B-1: the connectome of the significant cells; B-2: the changing trends of the significant cells. C) Cells indicating a U-shape relationship across the adult lifespan; C-1: the connectome of the significant cells; C-2: the changing trends of the significant cells. D) Cells showing an inverted U-shape relationship across the adult lifespan. D-1: the connectome of the significant cells; D-2: the changing trends of the significant cells.

### Comparisons between age-related SNC and FNC variations

After applied the well-matched structure-function template, five significant cells in the SNC matrix are matched with 11 significant cells in the FNC matrix as shown in Figure 5. Only two overlapping cells exhibited a similar changing trend of age-related SNC and FNC variations. One was the link between the middle occipital gyrus and insula, presenting a significant similarity between the two changing curves across the adult lifespan (*r*=0.53, *p*=1.23e-5). The other one was the link between the precuneus and cerebellum (*r*=0.28, *p*=0.029).

**Figure 5.**
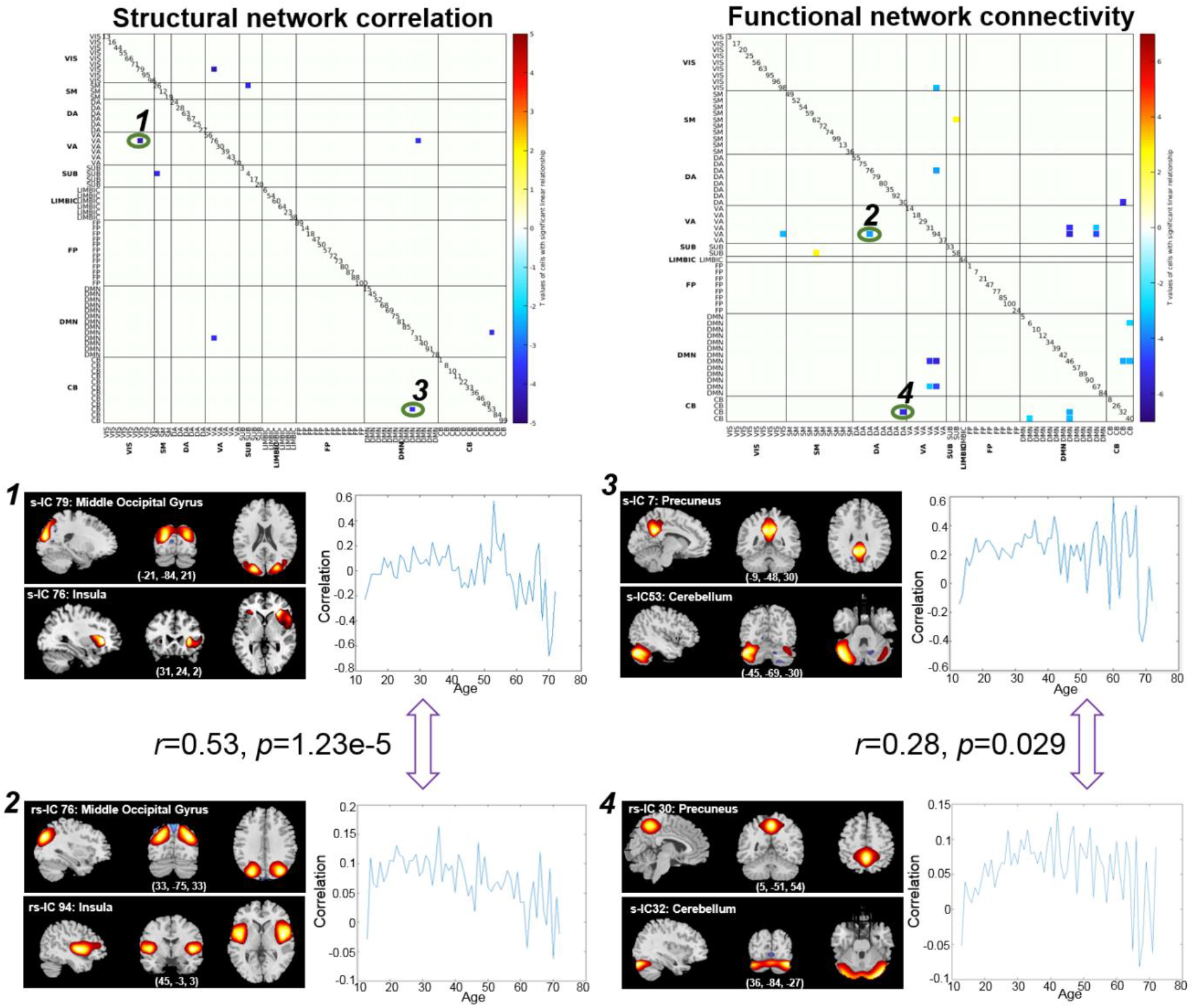
The cells presenting similar changing trends with age between SNC and FNC. Two cells (green circle) were revealed with significant similar changing trend: middle occipital gyrus-insula link (cell 1 and cell 2, *p* = 1.23E-5) and precuneus-cerebellum link (cell 3 and cell 4, *p* = 0.029).

## DISCUSSION

To the best of our knowledge, this is the first study to measure the age-related (co)variations of structural and functional connectivity/correlation using a very large sample (nearly 6000 subjects), based on the year-wise estimation from 13 to 72 years old, including both developmental and declining periods. Moreover, based on a well-matched structural-functional template, we estimated the changing trend with age and the turning age point of FNC and SNC carefully, and discovered that the turning age point of the U-shape curve is earlier for FNC than SNC. Our results indicated that 1) GMV exhibited a linearly declined relationship with age in most regions, while parahippocampus showed an inverted U-shape quadratic relationship with age; SNC presented a U-shape quadratic relationship with age within cerebellum, and inverted U-shape relationship primarily in the DMN and FP related connectivity. 2) FNC tended to linearly decrease within RSNs, especially in VIS and DMN. Early increase was revealed between RSNs, primarily in FP and DMN, which experienced decrease at older ages. U-shape relationship was also revealed to compensate for the cognition deficit in attention and SUB related connectivity at late years. 3) The connection between middle occipital gyrus and insula, as well as precuneus and cerebellum, exhibited similar age-related changing trends between SNC and FNC across the adult lifespan.

### Lifespan changes in gray matter volume

Most trajectories of the structural ICA-decomposed components showed a linearly declining relationship with age, primarily existed in the SM, DMN and FP network, consistent with previous studies (J. S. Allen et al., 2005; Liu et al., 2017; Spreng & Turner, 2013). For example, Spreng *et al.* also observed that GMV declined linearly with age in DMN across the adult lifespan of 18-96 years (Spreng & Turner, 2013). Allen *et al.* revealed a negative correlation between GMV and age in the frontal, parietal and visual networks (J. S. Allen et al., 2005). The parahippocampal, and hippocampus presented positive relationship with age, consistent with (Bagarinao et al., 2018). The increase may represent maturational changes, such as myelination, which would continue until late middle adulthood (He & Evans, 2014). Notably, the parahippocampus area further showed an inverted U-shape relationship, consistent with previous studies (J. S. Allen et al., 2005; Kalpouzos et al., 2009). An inverted U-shape relationship in the hippocampus was not revealed in this study, which appears inconsistent with previous observation that this region was vulnerable to aging and dementia. We did find a significant inverted U-shape relationship in s-IC23, with peaks at hippocampus and fusiform, however the partial R square for the quadratic relationship of s-IC23 was only 0.017. In this study we only reported the components which presented a partial R square value larger than 0.05. Moreover, selection bias in older participants in cross-sectional data is a potential limitation to the lower age-related reduction in the hippocampus of elder subjects as highlighted in (Nyberg et al., 2010). In addition, the GM volumes showed both positive and negative linear correlations with age in cerebellum and subcortical network. As shown in **Figure S2**, even though both s-IC49 and s-IC36 belongs to cerebellum network, the s-IC36 composed of the vermis and s-IC49 primarily composed of the posterior cerebellum. Previous studies have reported different volumetric trajectories in the anterior cerebellum/vermis, and posterior cerebellum. The anterior cerebellum and vermis follow a logarithmic pattern such that volume was largest in adolescents and dropped quickly during young adulthood (Bernard, Leopold, Calhoun, & Mittal, 2015). While the volumetric pattern of the posterior cerebellum seems to follow the protracted developmental pattern of the prefrontal cortex(Bernard et al., 2015). Moreover, it was recently suggested that this posterior cerebellar motor representation serves different functions than the anterior cerebellar motor representation (Donchin et al., 2012). Therefore, we would observe different relationship with age in the two areas. For the subcortical network, the s-IC20 presented negative relationship with age, with peaks at the caudate, while positive relationship was observed in s-IC4, which composed of thalamus. Previous studies have also reported different age-related volumetric variations in caudate (Walhovd et al., 2011) and thalamus (Bagarinao et al., 2018; Grieve, Clark, Williams, Peduto, & Gordon, 2005). As the thalamus is structurally connected to the hippocampus via mammillothalamic tract (Carlesimo, Lombardi, & Caltagirone, 2011) and functionally connected to the hippocampus as part of the extended hippocampal system (Stein et al., 2000), the volumetric trajectories of this area would be positive with aging similar to hippocampus.

### Lifespan changes in structural network connectivity

In order to make an extensive examination of age-related variations in both cortical properties and connectivity, we also studied how structural or functional connectivity changed with age across the adult lifespan. The DMN and FP related connectivity presented an inverted U-shape relationship, which are consistent with “last-in-first-out” rule: the late-maturing brain regions are revealed to be more sensitive to the deleterious effects of aging (Kalpouzos et al., 2009; Terribilli et al., 2011). The connectivity of these areas would mature after other brain areas, followed by atrophy, and then present a significant inverted U-shape tendency with age. Collin et al. also suggested that the dynamic changes in connectome organization throughout the lifespan follows an inverted U-shaped pattern (Collin & van den Heuvel, 2013). Moreover, since these brain regions are primarily involved in cognitive functions like attention, executive function and cognitive control, older brain will present increased activation in other related connectivity, for example, the dorsal_attention (DA) and ventral_attention (VA) network in this study, to reveal neural compensatory mechanisms (Rafael Romero-Garcia, Atienza, & Cantero, 2014), leading to a U-shape relationship. Besides, cerebellum also presented a U-shape relationship with age, which is consistent with previous studies (Brenhouse & Andersen, 2011; Durston et al., 2001). Notably, since the structural data were constructed based on the ICA-decomposed gray matter components, the age-related GMV variations could still reflect some SNC changes. For example, as shown in Figure 2A, the GMV linearly increased with age primarily in cerebellum and limbic system, while the other brain regions showed a linearly decreased relationship with age. Consistent with this, we also observed quadratic increased trends within the cerebellum network connectivity and quadratic decreased relationship in other brain network connectivity as shown in Figure 3B.

### Lifespan changes in functional network connectivity

Both linear and quadratic relationships with age were revealed in FNC cells. Early linearly increase in FNC were primarily observed between RSNs, which experienced decrease at older ages, for example the connectivity in STG, ACG, MCG and SFG areas, consistent with previously reported results (Betzel et al., 2014; Geerligs, Renken, Saliasi, Maurits, & Lorist, 2015). According to the “last-in-first-out” hypothesis revealed in the frontal and temporal areas as discussed above, the connectivity between frontal and temporal areas matured after other brain areas, followed by atrophy, and then exhibited an inverted U-shape tendency with age. The FNC within RSNs decreased linearly over the adult lifespan, especially in DMN and VIS, consistent with other studies (Andrews-Hanna et al., 2007; Geerligs et al., 2015; Tomasi & Volkow, 2012; L. Yan, Zhuo, Wang, & Wang, 2011). Late increase at older ages was observed in attention and SUB related area, which has been reported to compensate for the cognition deficit at late ages (Chen et al., 2018).

### Comparisons between changes in SNC and FNC

The relationships between age-related SNC changes and FNC variations are complex. Several previous studies reported that regions with few or no direct structural connections may exhibit high functional connectivity, which indicates presence of indirect connections between structure and function (Damoiseaux & Greicius, 2009; Honey et al., 2009). Kalinosky *et al.* further reported that the structural connectivity changes constructed using dMRI data were not related to the corresponding changes in FNC within RSNs (Kalinosky, Schindler-Ivens, & Schmit, 2013). In this study, we observed two matched SNC and FNC cells exhibiting a similar age-related changing trend across the adult lifespan as shown in Figure 5: 1) the connectivity between middle occipital gyrus and insula; 2) the connectivity between precuneus and cerebellum. The connectivity between middle occipital gyrus and insula was reported to be responsible for face emotion processing (Guo et al., 2015). The precuneus has also been identified to react to fearful faces (Zhao, Zhao, Zhang, Cui, & Fu, 2017) and is part of the extended face-processing network (Fox, Iaria, & Barton, 2009). Cerebellum was revealed to be implicated in rapidly coordinating information processing, aversive conditioning, and learning the precise timing of anticipatory responses (Auday, Taber-Thomas, & Perez-Edgar, 2018). Concerning the integration of contextual body signals in facial emotion perception, previous studies have shown an association with precuneus (contextual integration) and cerebellum (motor resonance) (Santamaría-García et al., 2019). Additionally, the turning point of quadratic relationship for age-related FNC variations was earlier than the turning point of SNC. FNC was revealed to detect brain activity through measuring variations related to blood flow, which are sensitive to environment changes (Deakin et al., 2008; Lahti, Holcomb, Medoff, & Tamminga, 1995). While SNC measures the links between pairs of covarying gray matter patterns, the deficits of which take time to manifest. These may partly help explain why age-related FNC variations would present an earlier changing point compare to age-related SNC changes.

### Limitations of the current study

There are several limitations in the current study. The first one is the lack of careful assessment of health status for the individuals included. As the data included a large sample size with a large age range, we believe the results may be more driven by the common characteristics of age-related changes. The second limitation is the cross-sectional nature of the data. Studies which used cross-sectional subjects may suffer from cohort effects and could not investigate changes over time within subjects compared to longitudinal studies. While longitudinal studies cannot totally replace cross-sectional studies for some limitations, such as the life expectancy of scanners (Salthouse, 2012). Third, although structural covariance of brain region volumes have been proved to be associated with both structural connectivity and transcriptomic similarity (R. Romero-Garcia et al., 2018; Yee et al., 2018), it is an indirect, group-wise measurement to scale the structural connectivity compared to tracking individual white matter fiber connectivity using diffusion magnetic resonance imaging (dMRI). Further work evaluating age-related variations using dMRI-based white matter connectivity is needed. Finally, the resolution of the fMRI images is sub-optimal (3.75×3.75×4.55mm) compared to the newer multi-band sequences (2-3 mm isotropic).

## ACKNOWLEDGMENTS

The authors thank Srinivas Rachakonda for lending his expertise on the GIFT toolbox functions, Helen Petropoulos for providing information on the fMRI data analyzed in this paper and the anonymous reviewers for their valuable comments and effort to improve the manuscript.

The authors report no biomedical financial interests or potential conflicts of interest.

## REFERENCES

Abe, O., Yamasue, H., Aoki, S., Suga, M., Yamada, H., Kasai, K., … Ohtomo, K. (2008). Aging in the CNS: Comparison of gray/white matter volume and diffusion tensor data. Neurobiology of Aging, 29(1), 102–116. doi:10.1016/j.neurobiolaging.2006.09.003

Abrol, A., Damaraju, E., Miller, R. L., Stephen, J. M., Claus, E. D., Mayer, A. R., & Calhoun, V. D. (2017). Replicability of time-varying connectivity patterns in large resting state fMRI samples. Neuroimage, 163, 160–176. doi:10.1016/j.neuroimage.2017.09.020

Allen, E. A., Erhardt, E. B., Damaraju, E., Gruner, W., Segall, J. M., Silva, R. F., … Calhoun, V. D. (2011). A baseline for the multivariate comparison of resting-state networks. Front Syst Neurosci, 5, 2. doi:10.3389/fnsys.2011.00002

Allen, J. S., Bruss, J., Brown, C. K., & Damasio, H. (2005). Normal neuroanatomical variation due to age: The major lobes and a parcellation of the temporal region. Neurobiology of Aging, 26(9), 1245–1260. doi:10.1016/j.neurobiolaging.2005.05.023

Andrews-Hanna, J. R., Snyder, A. Z., Vincent, J. L., Lustig, C., Head, D., Raichle, M. E., & Buckner, R. L. (2007). Disruption of large-scale brain systems in advanced aging. Neuron, 56(5), 924–935. doi:10.1016/j.neuron.2007.10.038

Ashburner, J., & Friston, K. J. (2005). Unified segmentation. Neuroimage, 26(3), 839–851. doi:10.1016/j.neuroimage.2005.02.018

Auday, E. S., Taber-Thomas, B. C., & Perez-Edgar, K. E. (2018). Neural correlates of attention bias to masked facial threat cues: Examining children at-risk for social anxiety disorder. Neuroimage Clin, 19, 202–212. doi:10.1016/j.nicl.2018.04.003

Bagarinao, E., Watanabe, H., Maesawa, S., Mori, D., Hara, K., Kawabata, K., … Sobue, G. (2018). An unbiased data-driven age-related structural brain parcellation for the identification of intrinsic brain volume changes over the adult lifespan. Neuroimage, 169, 134–144. doi:10.1016/j.neuroimage.2017.12.014

Bartsch, T., & Wulff, P. (2015). The Hippocampus in Aging and Disease: From Plasticity to Vulnerability. Neuroscience, 309, 1–16. doi:10.1016/j.neuroscience.2015.07.084

Bernard, J. A., Leopold, D. R., Calhoun, V. D., & Mittal, V. A. (2015). Regional cerebellar volume and cognitive function from adolescence to late middle age. Human Brain Mapping, 36(3), 1102–1120. doi:10.1002/hbm.22690

Betzel, R. F., Byrge, L., He, Y., Goni, J., Zuo, X. N., & Sporns, O. (2014). Changes in structural and functional connectivity among resting-state networks across the human lifespan. Neuroimage, 102, 345–357. doi:10.1016/j.neuroimage.2014.07.067

Brenhouse, H. C., & Andersen, S. L. (2011). Developmental trajectories during adolescence in males and females: A cross-species understanding of underlying brain changes. Neuroscience and Biobehavioral Reviews, 35(8), 1687–1703. doi:10.1016/j.neubiorev.2011.04.013

Burke, S. N., Gaynor, L. S., Barnes, C. A., Bauer, R. M., Bizon, J. L., Roberson, E. D., & Ryan, L. (2018). Shared Functions of Perirhinal and Parahippocampal Cortices: Implications for Cognitive Aging. Trends in Neurosciences, 41(6), 349–359. doi:10.1016/j.tins.2018.03.001

Calhoun, V. D., Adali, T., Pearlson, G. D., & Pekar, J. J. (2001). A method for making group inferences from functional MRI data using independent component analysis. Human Brain Mapping, 14(3), 140–151. doi:DOI 10.1002/hbm.1048

Calhoun, V. D., Adali, T., Pearlson, G. D., & Pekar, J. J. (2002). A method for making group inferences from functional MRI data using independent component analysis (vol 14, pg 140, 2001). Human Brain Mapping, 16(2), 131–131. doi:10.1002/hbm.10044

Cao, M., Huang, H., & He, Y. (2017). Developmental Connectomics from Infancy through Early Childhood. Trends in Neurosciences, 40(8), 494–506. doi:10.1016/j.tins.2017.06.003

Cao, M., Wang, J. H., Dai, Z. J., Cao, X. Y., Jiang, L. L., Fan, F. M., … He, Y. (2014). Topological organization of the human brain functional connectome across the lifespan. Dev Cogn Neurosci, 7, 76–93. doi:10.1016/j.dcn.2013.11.004

Carlesimo, G. A., Lombardi, M. G., & Caltagirone, C. (2011). Vascular thalamic amnesia: a reappraisal. Neuropsychologia, 49(5), 777–789. doi:10.1016/j.neuropsychologia.2011.01.026

Chen, Y., Zhao, X., Zhang, X., Liu, Y., Zhou, P., Ni, H., … Ming, D. (2018). Age-related early/late variations of functional connectivity across the human lifespan. Neuroradiology, 60(4), 403–412. doi:10.1007/s00234-017-1973-1

Collin, G., & van den Heuvel, M. P. (2013). The ontogeny of the human connectome: development and dynamic changes of brain connectivity across the life span. Neuroscientist, 19(6), 616–628. doi:10.1177/1073858413503712

Damoiseaux, J. S., & Greicius, M. D. (2009). Greater than the sum of its parts: a review of studies combining structural connectivity and resting-state functional connectivity. Brain Structure & Function, 213(6), 525–533. doi:10.1007/s00429-009-0208-6

Deakin, J. F., Lees, J., McKie, S., Hallak, J. E., Williams, S. R., & Dursun, S. M. (2008). Glutamate and the neural basis of the subjective effects of ketamine: a pharmaco-magnetic resonance imaging study. Arch Gen Psychiatry, 65(2), 154–164. doi:10.1001/archgenpsychiatry.2007.37

Donchin, O., Rabe, K., Diedrichsen, J., Lally, N., Schoch, B., Gizewski, E. R., & Timmann, D. (2012). Cerebellar regions involved in adaptation to force field and visuomotor perturbation. Journal of Neurophysiology, 107(1), 134–147. doi:10.1152/jn.00007.2011

Du, Y. H., & Fan, Y. (2013). Group information guided ICA for fMRI data analysis. Neuroimage, 69, 157–197. doi:10.1016/j.neuroimage.2012.11.008

Du, Y. H., Pearlson, G. D., Liu, J. Y., Sui, J., Yu, Q. B., He, H., … Calhoun, V. D. (2015). A group ICA based framework for evaluating resting fMRI markers when disease categories are unclear: application to schizophrenia, bipolar, and schizoaffective disorders. Neuroimage, 122, 272–280. doi:10.1016/j.neuroimage.2015.07.054

Durston, S., Hulshoff Pol, H. E., Casey, B. J., Giedd, J. N., Buitelaar, J. K., & Van Engeland, H. (2001). Anatomical MRI of the Developing Human Brain: What Have We Learned?. Journal of the American Academy of Child & Adolescent Psychiatry, 40(9), 1012–1020. doi: https://doi.org/10.1097/00004583-200109000-00009

Erhardt, E. B., Allen, E. A., Damaraju, E., & Calhoun, V. D. (2011). On network derivation, classification, and visualization: a response to Habeck and Moeller. Brain Connect, 1(2), 1–19.

Farokhian, F., Yang, C. L., Beheshti, I., Matsuda, H., & Wu, S. C. (2017). Age-Related Gray and White Matter Changes in Normal Adult Brains. Aging and Disease, 8(6), 899–909. doi:10.14336/Ad.2017.0502

Fjell, A. M., McEvoy, L., Holland, D., Dale, A. M., & Walhovd, K. B. (2014). What is normal in normal aging? Effects of aging, amyloid and Alzheimer’s disease on the cerebral cortex and the hippocampus. Progress in Neurobiology, 117, 20–40. doi: https://doi.org/10.1016/j.pneurobio.2014.02.004

Fjell, A. M., Westlye, L. T., Grydeland, H., Amlien, I., Espeseth, T., Reinvang, I., … Initiat, A. D. N. (2013). Critical ages in the life course of the adult brain: nonlinear subcortical aging. Neurobiology of Aging, 34(10), 2239–2247. doi:10.1016/j.neurobiolaging.2013.04.006

Fox, C. J., Iaria, G., & Barton, J. J. (2009). Defining the face processing network: optimization of the functional localizer in fMRI. Human Brain Mapping, 30(5), 1637–1651. doi:10.1002/hbm.20630

Freire, L., & Mangin, J. F. (2001). Motion correction algorithms may create spurious brain activations in the absence of subject motion. Neuroimage, 14(3), 709–722. doi:10.1006/nimg.2001.0869

Geerligs, L., Renken, R. J., Saliasi, E., Maurits, N. M., & Lorist, M. M. (2015). A Brain-Wide Study of Age-Related Changes in Functional Connectivity. Cerebral Cortex, 25(7), 1987–1999. doi:10.1093/cercor/bhu012

Grieve, S. M., Clark, C. R., Williams, L. M., Peduto, A. J., & Gordon, E. (2005). Preservation of limbic and paralimbic structures in aging. Human Brain Mapping, 25(4), 391–401. doi:10.1002/hbm.20115

Guo, W., Liu, F., Xiao, C., Zhang, Z., Liu, J., Yu, M., … Zhao, J. (2015). Decreased insular connectivity in drug-naive major depressive disorder at rest. Journal of Affective Disorders, 179, 31–37. doi: https://doi.org/10.1016/j.jad.2015.03.028

He, Y., & Evans, A. (2014). Magnetic resonance imaging of healthy and diseased brain networks. Frontiers in Human Neuroscience, 8, 890. doi:10.3389/fnhum.2014.00890

Honey, C. J., Sporns, O., Cammoun, L., Gigandet, X., Thiran, J. P., Meuli, R., & Hagmann, P. (2009). Predicting human resting-state functional connectivity from structural connectivity. Proceedings of the National Academy of Sciences of the United States of America, 106(6), 2035–2040. doi:10.1073/pnas.0811168106

Jafri, M. J., Pearlson, G. D., Stevens, M., & Calhoun, V. D. (2008). A method for functional network connectivity among spatially independent resting-state components in schizophrenia. Neuroimage, 39(4), 1666–1681. doi:10.1016/j.neuroimage.2007.11.001

Kalinosky, B. T., Schindler-Ivens, S., & Schmit, B. D. (2013). White matter structural connectivity is associated with sensorimotor function in stroke survivors. Neuroimage-Clinical, 2, 767–781. doi:10.1016/j.nicl.2013.05.009

Kalpouzos, G., Chetelat, G., Baron, J. C., Landeau, B., Mevel, K., Godeau, C., … Desgranges, B. (2009). Voxel-based mapping of brain gray matter volume and glucose metabolism profiles in normal aging. Neurobiology of Aging, 30(1), 112–124. doi:10.1016/j.neurobiolaging.2007.05.019

Kennedy, K. M., Erickson, K. I., Rodrigue, K. M., Voss, M. W., Colcombe, S. J., Kramer, A. F., … Raz, N. (2009). Age-related differences in regional brain volumes: a comparison of optimized voxel-based morphometry to manual volumetry. Neurobiology of Aging, 30(10), 1657–1676. doi:10.1016/j.neurobiolaging.2007.12.020

Lahti, A. C., Holcomb, H. H., Medoff, D. R., & Tamminga, C. A. (1995). Ketamine activates psychosis and alters limbic blood flow in schizophrenia. Neuroreport, 6(6), 869–872.

Liu, K., Yao, S. X., Chen, K. W., Zhang, J. C., Yao, L., Li, K., … Guo, X. J. (2017). Structural Brain Network Changes across the Adult Lifespan. Frontiers in Aging Neuroscience, 9. doi:ARTN 275 10.3389/fnagi.2017.00275

Meunier, D., Achard, S., Morcom, A., & Bullmore, E. (2009). Age-related changes in modular organization of human brain functional networks. Neuroimage, 44(3), 715–723. doi:10.1016/j.neuroimage.2008.09.062

Narvacan, K., Treit, S., Camicioli, R., Martin, W., & Beaulieu, C. (2017). Evolution of deep gray matter volume across the human lifespan. Human Brain Mapping, 38(8), 3771–3790. doi:10.1002/hbm.23604

Nyberg, L., Salami, A., Andersson, M., Eriksson, J., Kalpouzos, G., Kauppi, K., … Nilsson, L. G. (2010). Longitudinal evidence for diminished frontal cortex function in aging. Proc Natl Acad Sci U S A, 107(52), 22682–22686. doi:10.1073/pnas.1012651108

Power, J. D., Barnes, K. A., Snyder, A. Z., Schlaggar, B. L., & Petersen, S. E. (2012). Spurious but systematic correlations in functional connectivity MRI networks arise from subject motion. Neuroimage, 59(3), 2142–2154. doi:10.1016/j.neuroimage.2011.10.018

Raz, N., & Rodrigue, K. M. (2006). Differential aging of the brain: Patterns, cognitive correlates and modifiers. Neuroscience and Biobehavioral Reviews, 30(6), 730–748. doi:10.1016/j.neubiorev.2006.07.001

Romero-Garcia, R., Atienza, M., & Cantero, J. L. (2014). Predictors of coupling between structural and functional cortical networks in normal aging. Human Brain Mapping, 35(6), 2724–2740. doi:10.1002/hbm.22362

Romero-Garcia, R., Whitaker, K. J., Vasa, F., Seidlitz, J., Shinn, M., Fonagy, P., … Consortium, N. (2018). Structural covariance networks are coupled to expression of genes enriched in supragranular layers of the human cortex. Neuroimage, 171, 256–267. doi:10.1016/j.neuroimage.2017.12.060

Salthouse, T. A. (2012). Robust Cognitive Change. Journal of the International Neuropsychological Society, 18(4), 749–756. doi:10.1017/S1355617712000380

Santamaría-García, H., Ibáñez, A., Montaño, S., García, A. M., Patiño -Saenz, M., Idarraga, C., … Baez, S. (2019). Out of Context, Beyond the Face: Neuroanatomical Pathways of Emotional Face-Body Language Integration in Adolescent Offenders. Frontiers in behavioral neuroscience, 13, 34–34. doi:10.3389/fnbeh.2019.00034

Segall, J. M., Allen, E. A., Jung, R. E., Erhardt, E. B., Arja, S. K., Kiehl, K., & Calhoun, V. D. (2012). Correspondence between structure and function in the human brain at rest. Frontiers in Neuroinformatics, 6. doi:UNSP1010.3389/fninf.2012.00010

Silver, M., Montana, G., Nichols, T. E., & Alzheimer’s Disease Neuroimaging, I. (2011). False positives in neuroimaging genetics using voxel-based morphometry data. Neuroimage, 54(2), 992–1000. doi:10.1016/j.neuroimage.2010.08.049

Silver, M., Montana, G., Nichols, T. E., & Neuroimaging, A. D. (2011). False positives in neuroimaging genetics using voxel-based morphometry data. Neuroimage, 54(2), 992–1000. doi:10.1016/j.neuroimage.2010.08.049

Spreng, R. N., & Turner, G. R. (2013). Structural Covariance of the Default Network in Healthy and Pathological Aging. Journal of Neuroscience, 33(38), 15226–15234. doi:10.1523/Jneurosci.2261-13.2013

Stein, T., Moritz, C., Quigley, M., Cordes, D., Haughton, V., & Meyerand, E. (2000). Functional connectivity in the thalamus and hippocampus studied with functional MR imaging. AJNR Am J Neuroradiol, 21(8), 1397–1401.

Sui, J., He, H., Pearlson, G. D., Adali, T., Kiehl, K. A., Yu, Q., … Calhoun, V. D. (2013). Three-way (N-way) fusion of brain imaging data based on mCCA+jICA and its application to discriminating schizophrenia. Neuroimage, 66, 119–132. doi:10.1016/j.neuroimage.2012.10.051

Terribilli, D., Schaufelberger, M. S., Duran, F. L., Zanetti, M. V., Curiati, P. K., Menezes, P. R., … Busatto, G. F. (2011). Age-related gray matter volume changes in the brain during non-elderly adulthood. Neurobiology of Aging, 32(2), 354–368. doi:10.1016/j.neurobiolaging.2009.02.008

Toga, A. W., Thompson, P. M., Mori, S., Amunts, K., & Zilles, K. (2006). Towards multimodal atlases of the human brain. Nature Reviews Neuroscience, 7(12), 952–966. doi:10.1038/nrn2012

Tomasi, D., & Volkow, N. D. (2012). Aging and functional brain networks. Molecular Psychiatry, 17(5), 549–558. doi:10.1038/mp.2011.81

van den Heuvel, M. P., van Soelen, I. L., Stam, C. J., Kahn, R. S., Boomsma, D. I., & Hulshoff Pol, H. E. (2013). Genetic control of functional brain network efficiency in children. Eur Neuropsychopharmacol, 23(1), 19–23. doi:10.1016/j.euroneuro.2012.06.007

Vasa, F., Seidlitz, J., Romero-Garcia, R., Whitaker, K. J., Rosenthal, G., Vertes, P. E., … Consortium, N. (2018). Adolescent Tuning of Association Cortex in Human Structural Brain Networks. Cerebral Cortex, 28(1), 281–294. doi:10.1093/cercor/bhx249

Walhovd, K. B., Westlye, L. T., Amlien, I., Espeseth, T., Reinvang, I., Raz, N., … Fjell, A. M. (2011). Consistent neuroanatomical age-related volume differences across multiple samples. Neurobiology of Aging, 32(5), 916–932. doi:10.1016/j.neurobiolaging.2009.05.013

Wang, L. B., Su, L. F., Shen, H., & Hu, D. W. (2012). Decoding Lifespan Changes of the Human Brain Using Resting-State Functional Connectivity MRI. Plos One, 7(8). doi:ARTN e4453010.1371/journal.pone.0044530

Wu, K., Taki, Y., Sato, K., Kinomura, S., Goto, R., Okada, K., … Fukuda, H. (2012). Age-related changes in topological organization of structural brain networks in healthy individuals. Human Brain Mapping, 33(3), 552–568. doi:10.1002/hbm.21232

Xu, L., Groth, K. M., Pearlson, G., Schretlen, D. J., & Calhoun, V. D. (2009). Source-Based Morphometry: The Use of Independent Component Analysis to Identify Gray Matter Differences With Application to Schizophrenia. Human Brain Mapping, 30(3), 711–724. doi:10.1002/hbm.20540

Yan, C. G., Cheung, B., Kelly, C., Colcombe, S., Craddock, R. C., Di Martino, A., … Milham, M. P. (2013). A comprehensive assessment of regional variation in the impact of head micromovements on functional connectomics. Neuroimage, 76, 183–201. doi:10.1016/j.neuroimage.2013.03.004

Yan, L., Zhuo, Y., Wang, B., & Wang, D. J. (2011). Loss of Coherence of Low Frequency Fluctuations of BOLD FMRI in Visual Cortex of Healthy Aged Subjects. Open Neuroimag J, 5, 105–111. doi:10.2174/1874440001105010105

Yang, Z., Chang, C., Xu, T., Jiang, L., Handwerker, D. A., Castellanos, F. X., … Zuo, X. N. (2014). Connectivity trajectory across lifespan differentiates the precuneus from the default network. Neuroimage, 89, 45–56. doi:10.1016/j.neuroimage.2013.10.039

Yee, Y., Fernandes, D. J., French, L., Ellegood, J., Cahill, L. S., Vousden, D. A., … Lerch, J. P. (2018). Structural covariance of brain region volumes is associated with both structural connectivity and transcriptomic similarity. Neuroimage, 179, 357–372. doi:10.1016/j.neuroimage.2018.05.028

Yeo, B. T. T., Krienen, F. M., Sepulcre, J., Sabuncu, M. R., Lashkari, D., Hollinshead, M., … Buckner, R. L. (2011). The organization of the human cerebral cortex estimated by intrinsic functional connectivity. Journal of Neurophysiology, 106(3), 1125–1165. doi:10.1152/jn.00338.2011

Zhao, K., Zhao, J., Zhang, M., Cui, Q., & Fu, X. (2017). Neural Responses to Rapid Facial Expressions of Fear and Surprise. Front Psychol, 8, 761. doi:10.3389/fpsyg.2017.00761

Zuo, X. N., He, Y., Betzel, R. F., Colcombe, S., Sporns, O., & Milham, M. P. (2017). Human Connectomics across the Life Span. Trends in Cognitive Sciences, 21(1), 32–45. doi:10.1016/j.tics.2016.10.005

Zuo, X. N., Kelly, C., Di Martino, A., Mennes, M., Margulies, D. S., Bangaru, S., … Milham, M. P. (2010). Growing together and growing apart: regional and sex differences in the lifespan developmental trajectories of functional homotopy. Journal of Neuroscience, 30(45), 15034–15043. doi:10.1523/JNEUROSCI.2612-10.2010

